# Altered protein expression of galactose and *N*-acetylgalactosamine transferases in schizophrenia superior temporal gyrus

**DOI:** 10.1101/649996

**Authors:** Toni M. Mueller, Nikita R. Mallepalli, James H. Meador-Woodruff

## Abstract

Abnormal glycosylation in schizophrenia brain has been previously implicated in disruptions to protein trafficking, localization, and function. Glycosylation is an enzyme-mediated posttranslational modification and, along with reports of altered glycan biosynthesis and abnormal glycan levels in peripheral fluids, growing evidence for altered glycosylation pathway activity in schizophrenia prompted investigation of specific enzymes which may be responsible for these abnormalities. Glycosylation is a highly variable yet tightly controlled cellular process and, depending on the molecule(s) affected, altered glycan modifications can influence the trajectory of many key functional pathways. In a preliminary gene array study, abnormal transcript expression of 36 glycosylation-associated enzymes was identified in schizophrenia brain. Subsequently, protein expression abnormalities of enzymes which act on glucose (UGGT2), mannose (EDEM2), fucose (FUT8, POFUT2), and N-acetylglucosamine (MGAT4A, B3GNT8) have been reported in this illness. The current study investigates protein expression levels by western blot analysis of the galactose and N-acetylgalactosamine (GalNAc) transferases B3GALTL1, B4GALT1, C1GALT1, GALNT2, GALNT7, GALNT16, and GALNTL5, as well as a related chaperone protein, COSMC, in superior temporal gyrus of age- and sex-matched pairs of schizophrenia and non-psychiatrically ill comparison subjects (N = 16 pairs). In schizophrenia, reduced β-1,4-galactosyltransferase 1 (B4GALT1) and polypeptide GalNAc-transferase 16 (GALNT16) were identified. These findings add support to the hypothesis that dysregulated glycosylation-associated enzyme expression contributes to altered protein glycosylation and downstream substrate-specific defects in schizophrenia.

---

Past research on schizophrenia has identified abnormalities associated with multiple neurotransmitter pathways and cell signaling mechanisms. Protein post-translational modifications (PTMs) can affect the function, subcellular localization, interaction network(s), and transcriptional regulation of substrate proteins (Dantuma and van Attikum, 2016; Prabakaran et al., 2012; Ryšlavá et al., 2013). Glycosylation, which is the enzymatic attachment and modification of carbohydrate structures (called glycans), is a widespread PTM that has been previously implicated in schizophrenia pathophysiology (Bauer et al., 2010; Mueller et al., 2014; Stanta et al., 2010; Tucholski et al., 2013b, 2013a). Glycoproteins play important roles in regulating nearly all aspects of cellular organization, cell-cell communication, intracellular targeting and trafficking, protein secretion and uptake, cell signaling and developmental processes (Dennis et al., 2009; Hakomori and Igarashi, 1995; Hannun and Obeid, 2008; Moremen et al., 2012; Ohtsubo and Marth, 2006; Spiro, 2002; Varki, 2017; Varki et al., 2009). Given that defects of glycoprotein synthesis and processing can produce deleterious pleiotropic effects and many glycoprotein-mediated cellular functions are known to be perturbed in schizophrenia, we have hypothesized that abnormal expression of glycosylation-associated enzymes may underlie glycoprotein abnormalities in the illness and thereby contribute to multiple aspects of cellular dysfunction in schizophrenia.

Several neurotransmitter receptors, transporters, and associated molecules with atypical patterns of glycosylation have been identified in schizophrenia brain (Barbeau et al., 1995; Bauer et al., 2010; Gilabert-Juan et al., 2012; Mueller et al., 2014; Tucholski et al., 2013b, 2013a; Varea et al., 2012). Transcript-level studies have found mRNA levels of some glycosylation enzymes altered in this illness, and abnormal expression of specific glycan structures and monosaccharides in patient cerebrospinal fluid and blood serum have been reported (Ikemoto, 2014; Maguire et al., 1997; McAuley et al., 2012; Mueller et al., 2018; Narayan et al., 2009; Stanta et al., 2010). Previous glycobiology research in schizophrenia has focused on abnormal glycan expression and altered glycosylation of substrate proteins, while mechanisms which underlie glycosylation defects have not been as extensively explored. Recently, protein expression of glycosylation-associated enzymes has become a target of investigation. Glycosylation is an enzyme-mediated process and identification of specific enzymes that may be responsible for abnormal patterns of glycosylation in schizophrenia is a current gap in knowledge. A preliminary gene array study from our lab found abnormal mRNA expression of 36 key glycosylation enzymes clustered in 12 functional categories in this illness (Mueller et al., 2018). Altered protein levels of the glucosyltransferase UGGT2, the mannosidase EDEM2, N-acetylglucosamine (GlcNAc) transferases MGAT4A and B3GNT8, and fucosyltransferases FUT8 and POFUT2 have also been reported in schizophrenia (Kim et al., 2018; Kippe et al., 2015; Mueller et al., 2018). Although several galactose and/or N-acetylgalactosamine (Gal/GalNAc) modifying enzymes were implicated in our preliminary gene array study, the protein levels of these enzymes have not yet been investigated. Given that Gal/GalNAc is incorporated in nearly every subtype of glycan structure expressed in brain, abnormalities of enzymes which synthesize or modify Gal/GalNAc-containing glycans could contribute to previously identified alterations of protein *N*-glycosylation in schizophrenia. Guided by our transcript findings, we identified 7 Gal/GalNAc-modifying enzymes and 1 related enzyme chaperone that demonstrate detectable protein expression levels in postmortem human cortex: B3GALTL1, B4GALT1, C1GALT1, C1GALT1C1 (also called COSMC), GALNT2, GALNT7, GALNT16, and GALNTL5. We measured these targets in homogenates of superior temporal gyrus (STG; Brodmann area 22) from sex- and age-matched pairs of schizophrenia and non-psychiatrically ill comparison subjects.

## Materials and Methods

### Subjects and Tissue Acquisition

Samples of STG were obtained from the NIH NeuroBioBank at the Mt. Sinai School of Medicine, as described at https://neurobiobank.nih.gov/about-best-practices. Next of kin consent was obtained for each subject before being included in this collection. Subjects with schizophrenia were diagnosed using DSM-III-R criteria and brains were assessed micro- and macroscopically by board-certified neuropathologists. Inclusion criteria include diagnosis by at least two clinicians, documentation of psychosis before age forty, at least ten years of hospitalization for schizophrenia, and no signs of other neurodegenerative disorders (Powchik et al., 1993; Purohit et al., 1998). Neuropathological examination was conducted and subjects were excluded from this study if there was evidence of a history of drug/alcohol abuse, coma longer than six hours prior to death, or death by unnatural causes. Comparison subjects with no known history of psychiatric illness were obtained from the same collection. Sixteen pairs of schizophrenia and comparison subjects were matched for sex and age, and postmortem interval (PMI) when possible (Table 1). For each subject, 1 cm^3^ of the full thickness of grey matter from the left hemisphere of STG was dissected at autopsy and stored at −80°C until use.

**Table 1.**
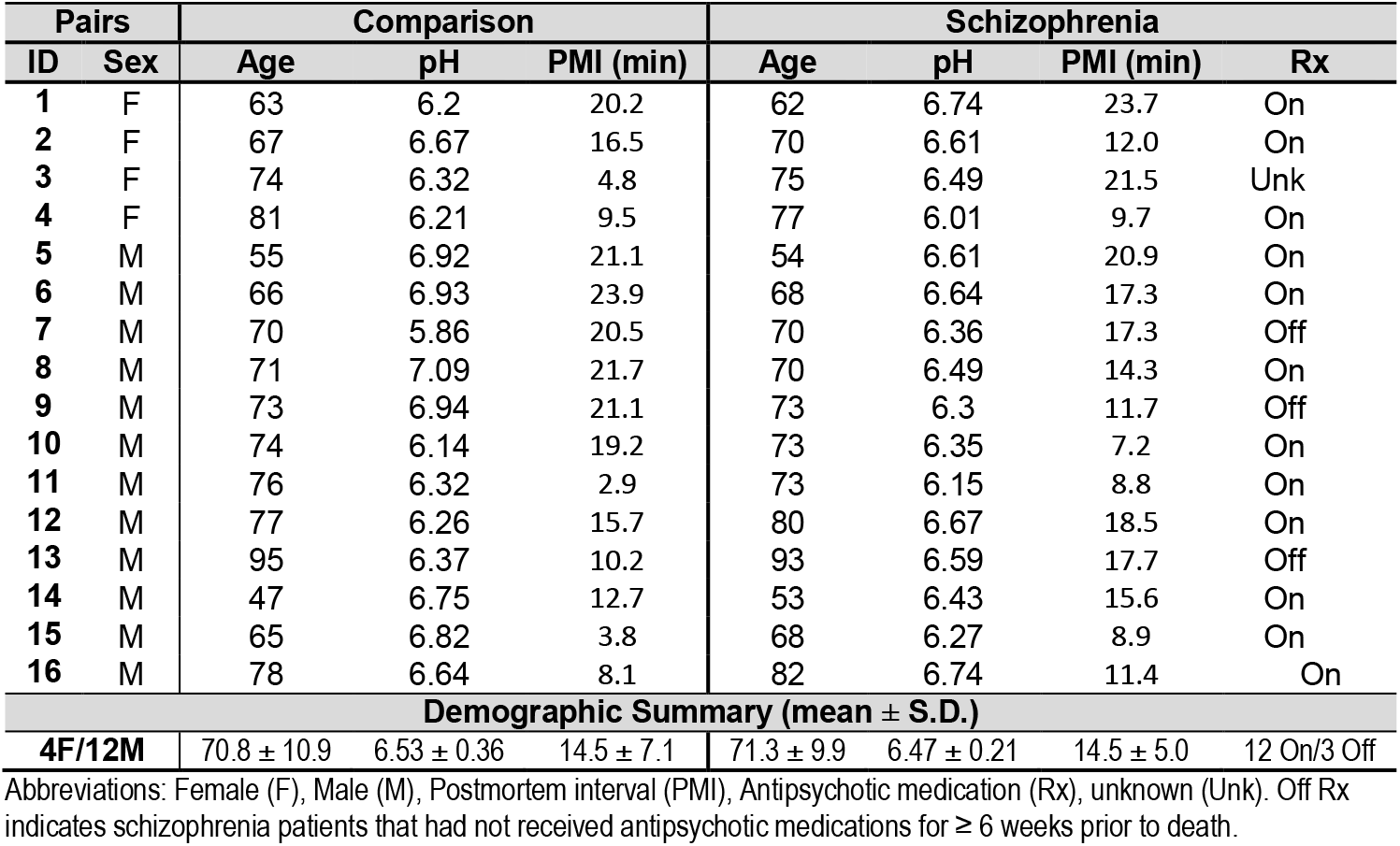
Paired Schizophrenia and Comparison Subject Demographic Information.

### Antipsychotic-Treated Rats

The Institutional Animal Care and Use Committee of the University of Alabama at Birmingham approved all animal studies. Male Sprague-Dawley rats were housed in pairs and treated with either 28.5 mg/kg of haloperidol decanoate (HALO, N = 10) or vehicle (CTRL, N = 10) via intramuscular injection every three weeks for a total of twelve injections over a 9 month period as previously described (Harte et al., 2005; Kashihara et al., 1986). Animals were euthanized by decapitation and brain tissue was immediately harvested. Samples of frontal cortex were dissected on wet ice, snap frozen on dry ice, and stored at −80°C until use.

### Sample Preparation

For both human and rat samples, tissue was homogenized with a Power Gen 125 homogenizer (Thermo Scientific, Waltham, MA) on wet ice in cold 5 mM Tris-HCL (pH 7.5) with 0.32 M sucrose and protease and phosphatase inhibitor tablets (Complete Mini, EDTA-free and PhosSTOP tablets; Roche Diagnostics, Indianapolis, IN). Protein concentrations were obtained for each homogenate using a BCA protein assay kit (Thermo Scientific) and samples were stored at −80°C until used for western blot analysis.

### Western Blot Analysis

Homogenized samples were thawed on wet ice and prepared with 6X loading buffer (4.5% sodium dodecyl sulfate (SDS), 0.02% bromophenol blue, 15% β-mercaptoethanol, and 36% glycerol in 170 mM Tris-HCL, pH 6.8) to a final 1x buffer concentration then heated to 70°C for 10 minutes and stored at −20°C until use. For each sample, 15μg of total protein was loaded in duplicate onto 4-12% Bis-Tris gel (Thermo Fisher) before being electrophoresed and transferred to 0.45 μM nitrocellulose membranes using a BioRad Semi-Dry Transblotter. A molecular mass standard was run on each gel (Novex Sharp Pre-stained Protein Standard, Life Technologies, Carlsbad, CA). Membranes were blocked for one hour at room temperature and probed with primary antibodies using conditions optimized to be within the linear range of detection for each primary antibody (Table 2). Valosin-containing protein (VCP) was used as an intralane loading control because its molecular weight is substantially different from the proteins of interest and has not previously shown alterations in schizophrenia (Bauer et al., 2009; Mueller et al., 2014; Stan et al., 2006). VCP antibodies from two different host species (mouse and rabbit) were used as needed to avoid cross-reactivity with the host species of primary antibodies for proteins of interest. Membranes were washed three times with Tris-buffered saline (TBS) containing 0.1% Tween-20 (TBST) then incubated with the appropriate IR-dye labeled secondary antibody in the same diluent as the primary antibody. The membranes were washed three times with TBST and once with MilliQ water prior to scanning on the Odyssey Infrared Imaging System (LI-COR Biosciences, Lincoln, NE). Image Studio Lite Version 5.0.2 (LI-COR Biosciences) was used to measure the signal of each target protein band at a resolution of 169 μm with the median right-left background signal intensity (3 pixels wide) subtracted. Expression levels were assessed similarly in HALO and CTRL rats for protein measures that were found significantly altered in schizophrenia.

**Table 2.** Antibody Conditions.

| Target Protein | Dilution | Buffer | Incubation | Manufacturer (Cat#) |
| --- | --- | --- | --- | --- |
| B3GALTL | 1:500 | 50% OBB | 16hr 4°C | Proteintech (14601-1-AP) |
| B4GALT1 | 1:500 | 50% OBB | 16hr 4°C | Abcam (ab121326) |
| GALNT2 | 1:500 | 50% OBB | 72hr 4°C | Abcam (ab140637) |
| GALNT7 | 1:500 | 50% OBB | 16hr 4°C | Proteintech (13962-1-AP) |
| GALNT16 | 1:1 000 | 2.5% Milk | 16hr 4°C | Abcam (ab168145) |
| GALNTL5 | 1:500 | Casein | 16hr 4°C | Proteintech (17131-1-AP) |
| C1GALT1 | 1:250 | 50% OBB | 16hr 4°C | Abcam (ab57492) |
| C1GALT1C1 | 1:500 | 50% OBB | 16hr 4°C | Proteintech (19254-1-AP) |
| VCP | 1:10 000 | 50% OBB | 1hr RT | Abcam (ab109240) |

All antibodies were obtained from Proteintech (Chicago, IL) or Abcam (Cambridge, MA). Buffers were diluted to the indicated concentration with Tris-buffered saline supplemented with 0.1% Tween-20. Abbreviations: Odyssey Blocking Buffer (OBB; #927-40100; LI-COR); non-fat dry milk (Milk); Casein Blocking Buffer (Casein; #927-40200; LI-COR); room temperature (RT).

### Data Analysis

The investigator who performed western blots was blind to subject diagnosis until study completion, and the investigator who performed statistical analyses was blind to diagnosis during data collection. Sample sizes were determined *a priori* using a power calculation (β = 0.2, π = 0.8) based on prior assessments of glycosylation enzyme protein expression in our lab (Kippe et al., 2015; Mueller et al., 2017). Data analysis was performed using Prism 6.07 (GraphPad Software Inc., La Jolla, CA) and STATISTICA 7.1 (StatSoft Inc., Tulsa, OK). Protein expression levels were calculated as the mean of the VCP-normalized target signal intensity. Data were assessed for statistical outliers using the ROUT method (robust regression and outlier removal) with Q = 1% (Motulsky and Brown, 2006) and then assessed for normal distribution using the D’Agostino and Pearson omnibus normality test. Between group differences were assessed by two-way paired Student’s t-tests for normally distributed data and by Wilcoxon matched pairs signed rank test for data that were not normally distributed. For significant dependent measures, *post hoc* assessments were performed including linear regressions of protein expression and subject age, tissue pH, and postmortem interval (PMI), and no significant associations with these continuous variables were identified. Due to small group sizes, *post hoc* Mann-Whitney tests were used to assess differences between Male/Female subjects. Since chronic antipsychotic administration is a potential confound of studies in elderly schizophrenia subjects, significantly different dependent measures were also assessed in haloperidol treated (N =10) or vehicle control (N = 10) Sprague-Dawley rats by two-way unpaired Student’s t-test following confirmation of normal distribution. For all statistical tests, α = 0.05.

## Results

### β-1,4-galactosyltransferase 1 (B4GALT1) and polypeptide *N*-acetylgalactosaminyltransferase 16 (GALNT16) are reduced in schizophrenia STG

The protein level of GALNT16 was reduced 44% (Figure 1 and Table 3; t(14) = 2.45, p = 0.028) and B4GALT1 was reduced 22% (Figure 2A and Table 3; W = −52, p = 0.043) in schizophrenia STG relative to matched comparison subjects. B4GALT1 protein expression was lower in female subjects (n = 6) relative to male subjects (n = 21) [t(24) = 2.30, p = 0.031]; however, given the small number of female subjects included in the study and the paired subject experimental design, it is unlikely that reduced B4GALT1 identified in schizophrenia is due to a sex effect on enzyme expression. No other enzymes measured in this study were found to be abnormally expressed in schizophrenia (Table 3).

**Figure 1.**
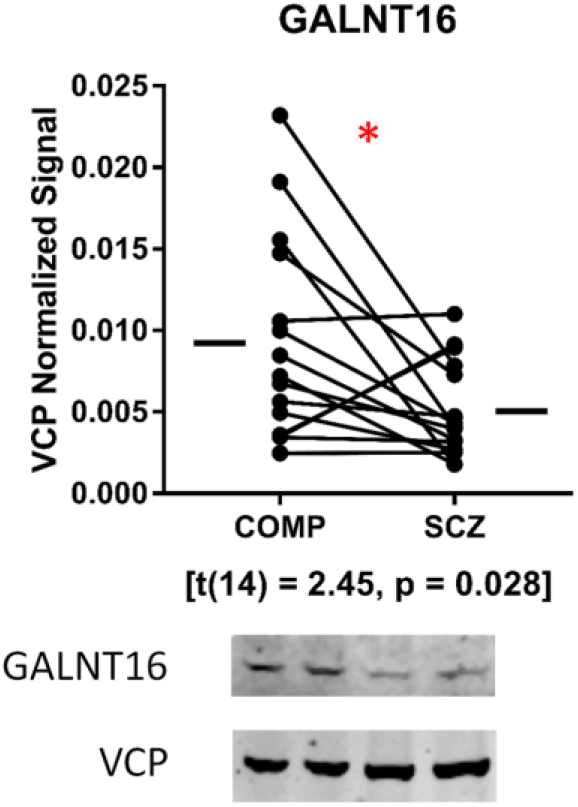
Expression of GALNT16 in STG of schizophrenia (SCZ) and comparison (COMP) subjects. Protein expression levels of GALNT16 are reduced 44% in schizophrenia STG. Data are presented as mean signal intensity of GALNT16 normalized to the signal intensity of intralane VCP for each subject. The mean for each diagnostic group is indicated by a horizontal bar. Images of representative western blots from a pair of age- and sex-matched subjects are shown. *p < 0.05

**Table 3.** Protein Expression.

| Table 3. Protein Expression |  |  |  |  |
| --- | --- | --- | --- | --- |
| Target Protein | Comparison | Schizophrenia | Test statistic | p-value |
| C1GALT1 | 0.060 ± 0.031 | 0.052 ± 0.022 | t (14) = 1.03 |  |
| C1GALT1C1 | 0.468 ± 0.225 | 0.400 ± 0.110 | t (13) = 0.29 |  |
| B3GALTL1 | 0.016 ± 0.008 | 0.020 ± 0.013 | W = 9 |  |
| B4GALT1 | 3.192 ± 2.019 | 2.502 ± 1.231 | W = -52 | 0.043 |
| GALNT2 | 0.124 ± 0.049 | 0.110 ± 0.032 | t (13) = 0.87 |  |
| GALNT7 | 0.018 ± 0.008 | 0.024 ± 0.016 | W = 5 |  |
| GALNT16 | 0.009 ± 0.006 | 0.005 ± 0.003 | t (14) = 2.45 | 0.028 |
| GALNTL5 | 0.065 ± 0.027 | 0.061 ± 0.017 | t (11) = 0.14 |  |

Values represent the mean VCP normalized signal intensity ± standard deviation for each diagnostic group.

### Chronic antipsychotic treatment does not alter the expression of B4GALT1

An animal model of chronic antipsychotic administration was tested to rule out the possibility that reduced protein levels of Gal/GalNAc transferases identified in schizophrenia may be attributed to long-term antipsychotic use. In rats chronically treated with haloperidol decanoate (HALO) versus vehicle controls (CTRL), no difference in the protein expression level of B4GALT1 was identified (Figure 2B) We were unable to detect GALNT16 protein expression in HALO and CTRL rats, most likely due to species-specific differences in the pattern of glycosylation enzyme protein expression, and therefore cannot completely exclude the possibility that antipsychotic treatment may influence protein expression of this enzyme.

**Figure 2.**
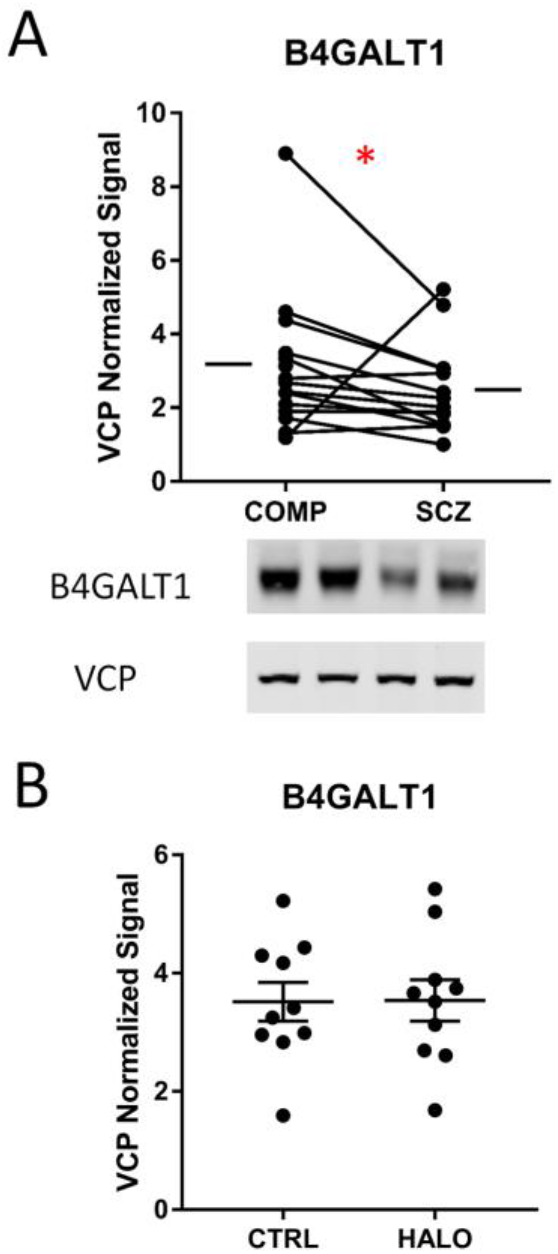
Expression of B4GALT1 in STG of schizophrenia (SCZ) and comparison (COMP) subjects. A) Protein expression levels of B4GALT1 are reduced 22% in schizophrenia STG. Data are presented as mean signal intensity of B4GALT1 normalized to the signal intensity of intralane VCP for each subject. The mean for each diagnostic group is indicated by a horizontal bar. Images of representative western blots from a pair of age- and sex-matched subjects are shown. *p < 0.05. B) There is no difference in the protein expression of B4GALT1 between rats chronically treated with either haloperidol (HALO) or vehicle (CTRL). Data are presented as the mean signal intensity of B4GALT1 normalized to the signal intensity of intralane VCP, with error bars indicating the S.E.M. for each treatment condition.

## Discussion

We measured 7 Gal/GalNAc transferases and 1 associated molecular chaperone, and found decreased expression of GALNT16 and B4GALT1 in schizophrenia STG. Each enzyme belongs to a unique subfamily of carbohydrate active enzymes grouped by sequence homology as well as shared structural and functional features of member proteins. In the Golgi apparatus, GALNT16 is one of 20 GALNT enzymes (Glycosyltransferase Family 27; GT27) that can initiate the first step of *O*-GalNAc glycan biosynthesis (Bennett et al., 2012), while B4GALT1 catalyzes the synthesis of lactose (a glucose-galactose disaccharide) and contributes to terminal galactosylation and the formation of lactose substructures within more complex glycoprotein glycans (Bydlinski et al., 2018). A subset of B4GALT1 is expressed in a secreted form following proteolytic cleavage, but it is unclear whether B4GALT1 exhibits extracellular enzymatic activity or merely serves as a cell-cell or cell-matrix recognition molecule (Bydlinski et al., 2018; Geisler et al., 2015; Yamaguchi and Fukuda, 2002).

The primary function of GALNTs is the initiation of *O*-GalNAc glycosylation, one of the most abundant types of glycosylation in animals (Bennett et al., 2012; Calvete and Sanz, 2008; Carraway and Hull, 1991). The *O*-GalNAc modification is highly expressed on mucins and proteins with mucin-like domains, and is often referred to as mucin-type *O*-glycosylation (Bennett et al., 2012; Brockhausen and Stanley, 2015; Carraway and Hull, 1991; Varki, 2017). Other forms of protein glycosylation are often initiated and regulated by the expression of one or two enzymes, whereas *O*-GalNAc structures can be initiated by any of 20 GALNT isoenzymes (GALNT1-20). Each GALNT isoenzyme has distinct patterns of substrate specificity, enzyme reaction kinetics, and variable patterns of cell-type specific expression levels over the course of development (Bennett et al., 2012; Brockhausen and Stanley, 2015; Iwasaki et al., 2003; Wandall et al., 1997). The GALNT enzyme family is one of the oldest and most evolutionarily conserved protein families and distinct yet partially overlapping substrate specificity of these 20 isoenzymes provides a mechanism for cellular compensation in the event that any individual GALNT is functionally impaired (Bennett et al., 2012). Many *O*-GalNAcylated substrates have been identified, but the substrate specificities of only a few GALNT isoenzymes have been thoroughly characterized (Bennett et al., 2012). Research has suggested a role for GALNT16 in the modulation of TGF-β signaling by interfering with Activin and BMP ligand-receptor binding (Bennett et al., 2012; Herr et al., 2008). These functional pathways are known to be perturbed in schizophrenia (Benes et al., 2007; Berretta, 2012; Kalkman, 2009; Karabon et al., 2015; Miller et al., 2011; Pietersen et al., 2014; Steiner et al., 2014). Thus, despite the possibility that other GALNT isoenzymes which were not assessed in the current study may be upregulated to partially compensate for reduced GALNT16 expression in schizophrenia, it is more likely that only a small subset of glycoproteins associated with TGF-β, Activin, and/or BMP signaling pathways may be impacted by reduced levels of GALNT16.

In our study, we found GALNT16 decreased in schizophrenia, but we did not identify expression differences for enzymes involved in subsequent steps of *O*-GalNAc extension or another GALNT enzyme from the same GT27 subfamily (GT27-Ib), GALNT2. Glycan structures initiated by GALNTs can be subsequently modified to produce one of 4 core structures or, if not further modified, the GalNAc-Ser/Thr motif called the Tn-antigen (Brockhausen and Stanley, 2015). C1GALT1 (Core 1 β-1,3-galactosyltransferase, also called T-synthase) and its chaperone C1GALT1C1 (C1GALT1-specific chaperone, also called COSMC) which act downstream of GALNT1-20 to produce a Core 1 *O*-GalNAc structure (T-antigen), were both normally expressed in schizophrenia. Another enzyme immediately downstream of GALNT1-20 is ST6GAL1 (β-galactoside α-2,6-sialyltransferase 1), which is necessary for the synthesis of the sialyl Tn-antigen. Both ST6GAL1 upregulation and enhanced expression of the sialyl TN-antigen are well-known biomarkers for several types of cancer (Li and Ding, 2018). ST6GAL1 protein levels are also not different in schizophrenia STG (unpublished results). Together, these data suggest that reduced GALNT16 levels could potentially impair the formation of multiple *O*-GalNAc structures on molecules associated with TGFβ signaling in schizophrenia.

The B4GALT enzyme family (Glycosyltransferase Family 7; GT7) is similarly composed of multiple isoenzymes, however only 7 B4GALTs are expressed in humans, each with variable affinity for glycoprotein, glycolipid, and/or proteoglycan substrates (Bydlinski et al., 2018; Lee et al., 2001). B4GALT1, which we found reduced in schizophrenia, is a key mediator of *N*-glycan galactosylation (Bydlinski et al., 2018). Although B4GALT2-4 can partially compensate for defects in B4GALT1 expression, the prevalence of immature *N*-glycans is increased when B4GALT1 is knocked out *in vitro* (Bydlinski et al., 2018). Interestingly, the microRNA miR-124-3p is an upstream regulator of B4GALT1 expression, and higher levels of miR-124-3p are associated with inhibition of B4GALT1 (Liu et al., 2016). A composite feed-forward loop composed of miR-124-3p, the transcription factor EGR1 (early growth response gene 1), and the target gene SKIL (SKI-like protooncogene) has been identified in schizophrenia and upregulation of miR-124-3p has been reported in drug-free schizophrenia patients (Xu et al., 2016).

Increased expression of Gal/GalNAc associated enzymes and glycan structures are common in various cancers (Schneider et al., 2017; Xie et al., 2018, 2016; Zhou et al., 2013, 2012) and schizophrenia patients diagnosed with cancer tend to have a longer delay to diagnosis, worse treatment outcomes, and higher mortality rates (Abdullah et al., 2015; Bergamo et al., 2014; Irwin et al., 2017; Ishikawa et al., 2016; Lawrence et al., 2000; Safdieh et al., 2017; Sathianathen et al., 2019; Zhuo et al., 2017). Since we have identified decreased levels of B4GALT1 and GALNT16 in schizophrenia, it is possible that established biomarkers for cancer diagnosis do not have the same degree of predictive accuracy for schizophrenia patients. While cancer incidence and outcomes in schizophrenia could also be due to lifestyle or other factors, poor survival rates may also reflect a physiological difference in biomarker expression that could contribute to delays in cancer diagnosis and/or reduced response to treatment. Future research on cancer risk and treatment outcomes in patients with major mental illness should examine differences related to glycosylation-associated biomarkers to shed light on molecular features of comorbid schizophrenia and cancer diagnoses.

As with most postmortem studies, some limitations must be considered in the interpretation of our findings. We sought to minimize confounds related to a lifetime of chronic antipsychotic use by testing significantly different dependent measures in a rodent model of chronic haloperidol administration. Unfortunately, GALNT16 was below the threshold for detection by western blot analysis in our rodent model, which we believe to represent a species-specific difference in the pattern of GALNT isoenzymes expressed in adult rodent brain. We did not identify a difference in B4GALT1 levels in haloperidol-treated rodents versus controls.

We found protein expression differences for two important Gal/GalNAc transferases, GALNT16 and B4GALT1. Increased levels of these enzymes and/or their downstream effectors have been implicated in relation to various forms of cancer and are, for some cancers, established biomarkers for diagnostic and prognostic screening. It is possible that reduced levels of these enzymes, as we found in schizophrenia, may indicate a mechanism that confers reduced risk for some forms of cancer or, alternatively, that established glycosylation-associated biomarkers used to detect cancer may be less effective in the schizophrenia population. Abnormal *N*-glycan galactosylation by B4GALT1 could also contribute to N-glycosylation abnormalities of neurotransmission associated molecules and downstream alterations of substrate trafficking and function suggested in this illness.

## Author Disclosures

### Contributors

Authors TMM and JHM-W designed the study. Author NRM executed experimental protocols. Author TMM managed literature searches and performed statistical analyses. Authors TMM and NRM collected data and wrote the first draft of the manuscript. All authors contributed to and have approved the final manuscript.

## Acknowledgements

The authors gratefully acknowledge Dr. Rosalinda Roberts and the Alabama Brain Collection for postmortem cortical samples used for antibody optimization. We also thank Chase Ryan-Embry for assistance with preliminary experiments and antibody optimization. Author TMM would like to additionally acknowledge the important role of Dr. W. Mueller, Ms. PRF Mueller, Mr. G. Leban, and Mr. O. Leban for providing moral support, encouragement, and constructive feedback during the entirety of the project.

